# A universal fluorescence-based toolkit for real-time quantification of DNA and RNA nuclease activity

**DOI:** 10.1101/548628

**Authors:** Emily C. Sheppard, Sally Rogers, Nicholas J. Harmer, Richard Chahwan

**Author notes:** Authors contributed equally to this work (Emily C. Sheppard, Sally Rogers). **Corresponding author:** Richard Chahwan.

## Abstract

DNA and RNA nucleases play a critical role in a growing number of cellular processes ranging from DNA repair to immune surveillance. Nevertheless, many nucleases have unknown or poorly characterized activities. Elucidating nuclease substrate specificities and co-factors can support a more definitive understanding of cellular mechanisms in physiology and disease. Using fluorescence-based methods, we present a quick, safe, cost-effective, and real-time versatile nuclease assay, which uniquely studies nuclease enzyme kinetics. In conjunction with a substrate library we can now analyse nuclease catalytic rates, directionality, and substrate preferences. The assay is sensitive enough to detect kinetics of repair enzymes when confronted with DNA mismatches or DNA methylation sites. We have also extended our analysis to study the kinetics of human single-strand DNA nuclease TREX2, DNA polymerases, RNA, and RNA:DNA nucleases. These nucleases are involved in DNA repair, immune regulation, and have been associated with various diseases, including cancer and immune disorders.

## Introduction

DNA and RNA nucleases are a hallmark of a growing number of cell signalling cascades including the DNA damage response and immune diversification. They dictate many DNA repair pathway choices by controlling the DNA damage substrates created that dictate downstream options during the signalling cascade. The complex roles DNA and RNA nucleases play in DNA repair pathways underpin several premature ageing-, immune-, and tumour-related syndromes. All of which can result from aberrations in the structural and/or catalytic functions of DNA and RNA nucleases (reviewed in (1) and (2)). Despite their importance, the activities of numerous human DNA nucleases are still debated (e.g. Mre11, CTIP, etc. (3, 4)). There are also several yet uncharacterised proteins harbouring predicted nuclease domains in mammalian genomes (5–8).

Even with their myriad and complex functions, DNA/RNA nucleases can be broadly defined by their substrate specificity, directionality of resection, and processivity. A significant restrictive limitation in nuclease studies has been the adequate identification of their catalytic functions and/or their relative activities against differing DNA/RNA intermediates. Conventional nuclease assays predominantly involve the use of radioactive labelling to visualise DNA substrates on an agarose gel (9–11). The use of radioactive isotopes delivers highly specific, sensitive assays that are free from interference. However, these assays are often inefficient, time-consuming, qualitative, and potentially hazardous (12, 13). Additionally, the assays are discontinuous, and must be stopped at discreet, often arbitrary, time points before measuring readouts (14). Whilst this can provide an indication of reaction rate, it does not allow for real-time visualisation of the catalytic resection activity.

Radiolabelled oligonucleotides are gradually being replaced with fluorescent nucleic acid stains such as DAPI (15) and other commercially-available dyes including, but not limited to, Midori Green, SYBR Green I and Acridine Orange (16). PicoGreen (PG) is a commercially available dye that emits a fluorescent signal upon intercalation with double-stranded DNA (dsDNA), emitting a fluorescent signal 1,000-fold stronger compared to when it is free in solution. In comparison, the Quant-It (QI) mircoRNA assay dye can specifically recognise short RNAs (<40-nts) and other nucleic acids. The superior sensitivity can quantify picogram or nanogram levels of dsDNA, respectively, unlike other fluorescent dyes, including Hoechst (17), ethidium bromide (17), EvaGreen (18), SYBR Green (18) and YOYO-1 (18). Several other sophisticated fluorescent techniques have been devised, building upon the use of FRET, whereby fluorescence is either quenched or dequenched following nuclease activity (19). Methods involving graphene oxide surfaces (20), electrochemical redox reactions (21–23), complexing of DNA with a polycationic polymer (24), or immobilising nucleotides on magnetic beads (25) have also been developed. While highly sensitive, these methods have been designed purely for the detection of a very limited number of DNA and RNA nucleases rather than for their characterisation (26).

Similar to other nucleic acid dyes, PG has proven to be a versatile DNA stain in different experimental conditions. PG has been used to visualise dsDNA in agarose electrophoresis as a quality control marker to identify fragmented and nicked DNA (27). It has also been implemented in flow cytometry analysis of cell-free DNA which can increase in certain pathologies, such as cancer and autoimmune syndromes (28). PG exhibits readily detectable, albeit reduced, fluorescence readings at temperatures of 37°C, despite manufacturer recommendations to work at room temperature (29). As PG very preferentially binds dsDNA compared to ssDNA and RNA structures (30), it has been an important feature of discontinuous enzyme-mediated DNA-modifying studies, such as nucleases (31), helicases (32), polymerases (33, 34), polymerase inhibitors (35), telomerases (36), and primases (37). However, the full potential of PG and QI in real-time visualisation of reaction progression remains limited (38).

Here, we propose a PG and QI fluorescence-based toolkit for performing real-time fluorescent nuclease assays. Our toolkit has been optimised for a range of important representative nucleases, including: DNase I (a non-specific nuclease), T7 exonuclease (a 5’-3’ bacteriophage nuclease) (14), Exonuclease III (a 3’-5’ *E. coli* nuclease) (39), human Trex2 (a 3’-5’ ssDNA nuclease) (40), a viral nickase, Nt.CviPII (41), RNase A (a ssRNA nuclease) and RNase H (targets RNA in RNA:DNA hybrids). Furthermore, we designed a library of nucleotide substrates to account for these enzymes’ differential activities. Our DNA substrates are designed with biotin-TEG-modified 3’ or 5’ termini to provide insight into enzyme directionality. The addition of streptavidin to the biotinylated ends protects the substrate from resection at the modified end, as was demonstrated in a previous resection assay (10). This oligonucleotide library can be used to study a multitude of uncharacterised nucleases, and their substrate preferences, to elucidate their roles in DNA repair and genomic maintenance. To extend the power of our approach, we have designed physiologically relevant substrates containing mismatches and methyl-cytosines. It has been posited that repair nucleases resect along a methylated sequence of DNA, thus permanently removing epigenetic markers. Upon recognition of a mismatch, the mismatch repair (MMR) machinery generates incisions either side of the error, allowing the 5’-3’ exonuclease Exo I to resect through the mismatch, permitting polymerases to accurately replicate across the resulting ssDNA tract (42). As such, it would be interesting to determine whether this mismatch is permissive to nucleases in general.

Our assay relies on the loss or gain of the DNA duplex structure, and so is potentially useful for study of other enzymes that reconstitute or compromise the DNA-pairing structure. As an example, we demonstrate the observation of polymerase activity by monitoring an increase in fluorescence as the complementary strand is synthesised by the nuclease-deficient Klenow fragment DNA polymerase. PG has previously been used to visualise polymerase activity, but to our knowledge, this is the first time it has been described in a real-time, continuous assay (33). Our work provides a robust and versatile universal toolkit to characterise both DNA and RNA nucleases, and determine their substrate preferences with high resolution and sensitivity. This assay can be adapted and modified to suit a wide range of DNA repair applications. This fluorescence-based method is faster, cheaper, and safer than conventional DNA/RNA resection assays using radioisotopes. Additionally, it works cross-species and is platform agnostic. Furthermore, this protocol is sensitive enough for enzyme reaction kinetic calculations and can distinguish the structural preferences exhibited by an enzyme for its substrate.

## Materials and Methods

### Oligonucleotides

DNA substrates were prepared by diluting plasmid DNA in HyClone water™ (GE healthcare), and unmodified HPLC-purified oligonucleotide substrates (Eurofins) and modified HPLC-purified oligonucleotides (Integrated DNA Technologies) in 1X Annealing Buffer (Sigma-Aldrich) for DNA substrates and siMAX™ dilution buffer (Eurofins) for all RNA substrates. Oligonucleotides were designed and optimised against secondary structure formation using the ‘Predict a Secondary Structure Web Server’ (https://rna.urmc.rochester.edu/RNAstructureWeb/Servers/Predict1/Predict1.html)(43) and annealed at a 1:1 molar ratio. Table 1 shows all the oligonucleotides and their respective illustrations, while Table S1 lists the olignonucleotide sequences and modifications.

**Table 1:**
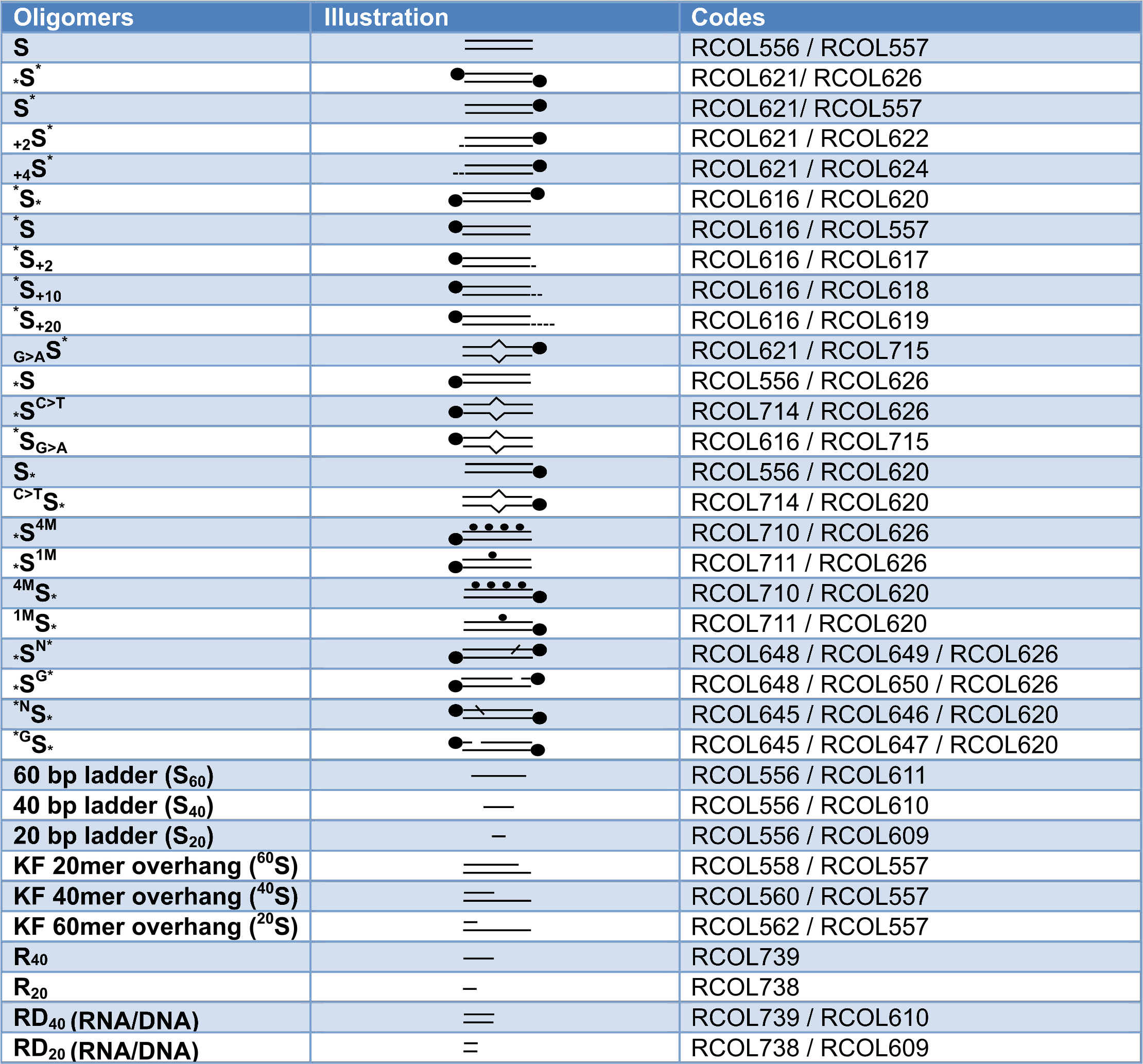
Oligonucleotides and their respective stick and ball illustrations

### Nucleases and PG preparation

The nucleases used were RQ1 RNase-Free DNase I (Promega), T7 exonuclease (New England Biolabs), Exonuclease III (New England Biolabs), Trex2 (Stratech), Klenow Fragment (3’ → 5’ exo-; New England Biolabs), RNase A (Thermo Fisher) and RNase H (New England Biolabs). In preparation for each assay, nucleases used were diluted on ice in their appropriate storage buffers, omitting glycerol. DNase I, T7 exonuclease, Exonuclease III, Trex2 and Klenow fragment storage buffers and reaction buffers were all filtered prior to use, and autoclaved where possible.

The PG reagent from the Quant-iT™ PicoGreen™ dsDNA Assay Kit (Invitrogen) was prepared immediately before use by making a 1:200 dilution of the PG in TE buffer (10 mM Tris-HCl, 1 mM EDTA, pH 7.5) and 40% (v/v) glycerol. For the RNase assays, the dye from the Quant-iT™ microRNA Assay Kit (Invitrogen) was prepared by diluting the microRNA reagent A into buffer B in a 1/2000 dilution, as detailed in the protocol for the kit.

### Experimental procedure

Each DNA substrate reaction mixture contained 50 nM DNA substrate, 1X reaction buffer (specific for each enzyme), 50 μL PG solution, 0.02 mg/mL streptavidin (cat. 21125, Thermo Fisher Scientific) if required, 0.25 mM dNTPs if required, 5 μL enzyme or relevant storage buffer. For the DNA nucleases, Milli-Q water was added to bring the total reaction volume to 100 μL. The RNA nuclease reaction mixtures, contained 50 nM RNA substrate 90 μL Quant-iT reagent and 5 μL enzyme or relevant storage buffer. RNase A does not require a specific reaction buffer for its activity, and therefore the enzyme was instead diluted in TE buffer. RNase-free HyClone™ Water (cat. SH305380, Fisher Scientific) was used to bring the reaction volume to 100 μL,. Samples were tested in a 96-well, black flat bottom plate (cat. M9685, Sigma-Aldrich). The final components added were the storage buffers, then the enzyme mixtures, in order to start the reaction.

All storage buffers and reaction buffers were made according to the recipes available on their respective NEB and Thermo Fisher web pages.

A CLARIOstar microplate reader (BMG labtech) was pre-heated to 37°C. Samples were read every 40-50 s for 30-60 mins. Excitation and emission wavelengths used were 483-15 nm and 530-30 nm, with a focal height of 10.2, 20 flashes per well, with a shake before each read.

### Data analysis

For statistical analysis of the data, one-way ANOVA with the Tukey’s post-hoc tests were used. This was implemented using GraphPad Prism v7.03. An example of the workflow is available in Supplementary Materials.

## Results

### Establishing assay parameters using DNase I

The foundation of the nuclease assay reported here is to monitor the corresponding loss of the fluorescent signal as dsDNA is degraded in response to DNA metabolizing enzymes, primarily DNA nuclease activities. As illustrated (Fig.1a), PG binds to dsDNA and produces a fluorescent signal that is approximately 400-fold more intense compared to when it is free in solution. Upon the addition of a nuclease, such as DNase I, the duplex structure is lost as the enzyme digests the DNA.

A calibration curve was established to determine the linear range in which the concentration of DNA is directly proportional to the fluorescent signal (Fig. 1b). We compared both plasmid DNA (13.1 kb) and oligonucleotide DNA (80 bp) and determined that the maximum concentration of DNA that could be used in the assay was 2.5 ng/μL regardless of the length and structure of the DNA. Above 2.5 ng/μL this, the fluorescent signal plateaued irrespective of the increase in DNA concentration (Fig. 1b).

**Figure 1:**
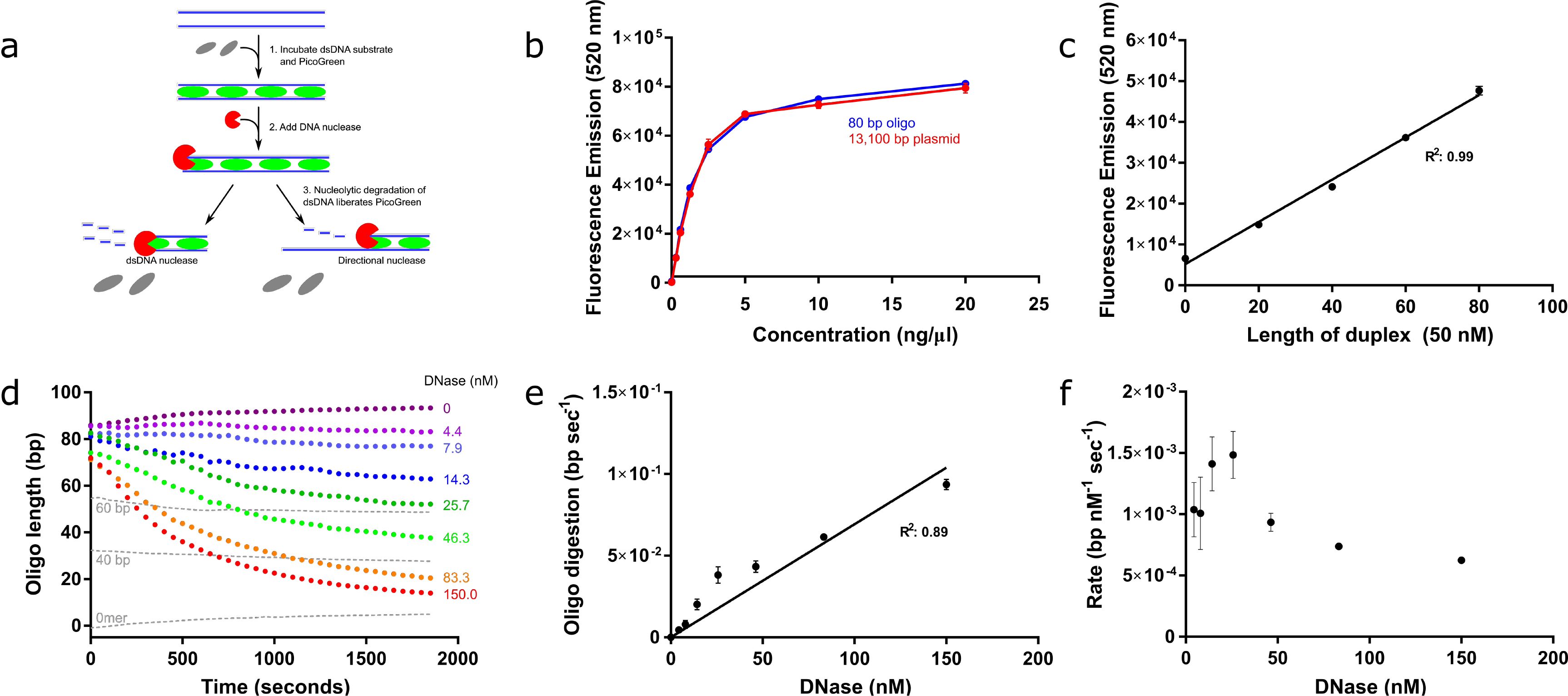
Optimisation of the fluorescence signal and assay parameters. **a**, Schematic depicting the method by which PG (green) intercalates with dsDNA (blue) to emit a fluorescent signal. Digestion of this PG:DNA complex by nucleases (red) disrupts the duplex structure and causes a concomitant reduction (grey) in signal that can be measured in real-time. **b,** Calibration curve indicating the fluorescent signal emitted by increasing concentrations of 80 bp dsDNA and 13.1 kb plasmid DNA. **c,** Standard curve composed of 80, 60, 40 and 20 bp sequences. The point shown at 0 on the x-axis is an 80-nt ssDNA oligomer to represent the end product of complete resection. **d,** DNase I titration on an 80 bp dsDNA substrate. Dotted grey lines represent controls containing standard duplexes of intermediate sizes. **e,** Extracted maximum gradient from (d) to determine the reaction rate at increasing concentrations of DNase I. **f,** Maximum rate analysis of resection per nM DNase I per second. Error bars represent SEM; n=3 in all cases.

As an alternative to converting fluorescence to molar concentrations, which vary according to the length of the DNA substrate (Fig. S1a-d), we generated a standard curve based on the signal produced from 80-, 60-, 40- and 20-bp substrates, as well as a 0-bp single-stranded 80-mer oligomer to represent the final resection product of other resection nucleases (Fig.1c). PG binds ssDNA with a lower affinity than dsDNA, as indicated by the standard curve. For resection nucleases (Fig. 2 and 3) as opposed to DNase, this is a more appropriate end-point of the reaction rather than the comparatively minor fluorescent signal produced in the absence of substrate. This standard curve shows a linear increase in fluorescence with the length of duplex DNA (R^2^ = 0.99).

**Figure 2:**
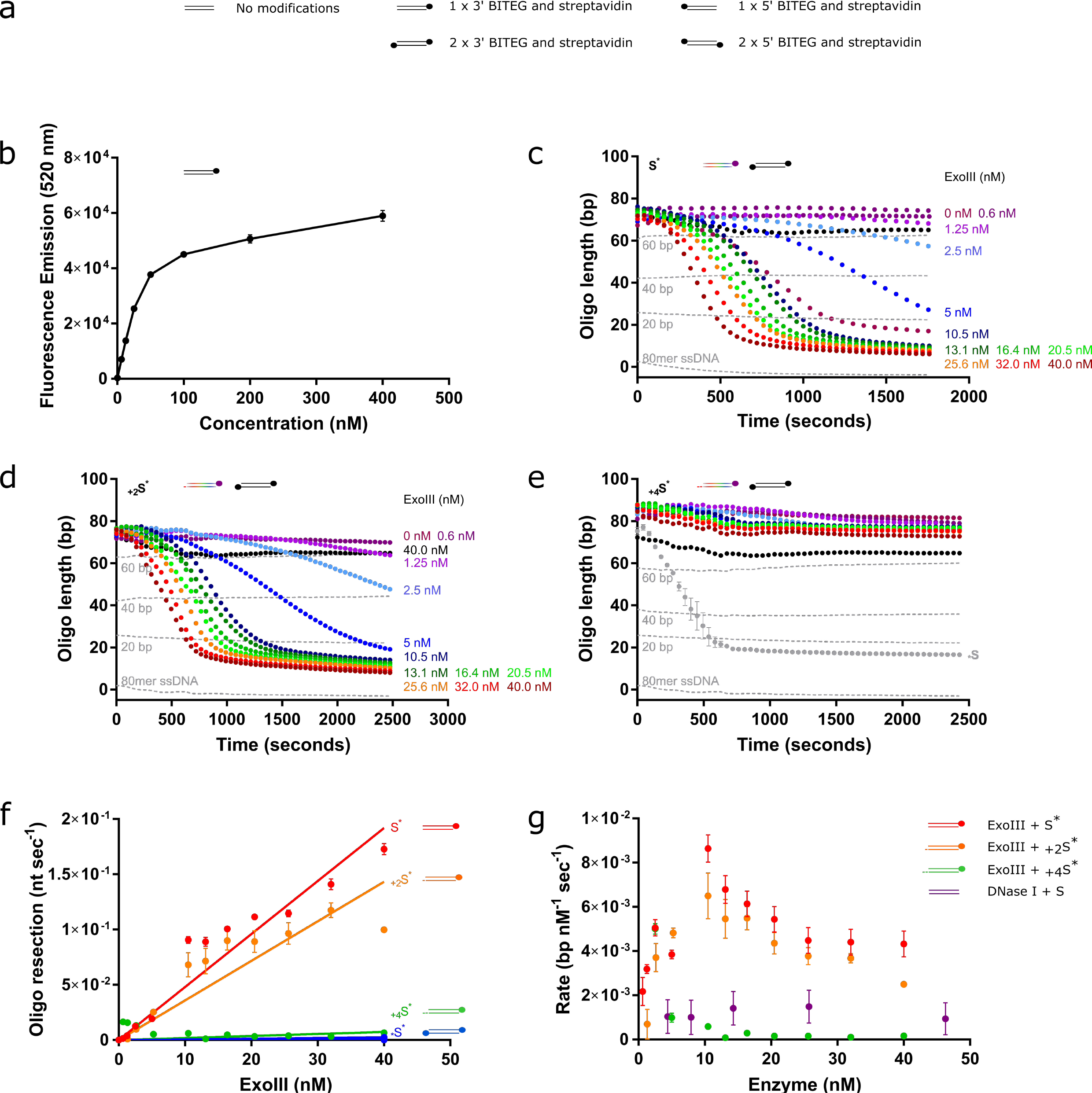
Assay validates inhibitory effect of increasing lengths of a 3’ overhang on 3’-5’ nuclease, ExoIII. **a**, Key to stick and ball illustrations of the substrates. The parallel horizontal lines (=) represent the dsDNA substrate, and the coloured-in circle represents the combination of both biotin and streptavidin (●). **b**, Calibration curve depicting the fluorescent signal according to increasing concentrations of 80-bp oligomer substrate with one terminal BITEG modification and in the presence of 0.02 mg/ml streptavidin. **c-e**, ExoIII titrations as depicted in the figures on 50 nM substrates presenting a blunt terminus (c), 2-nt (d) or 4-nt (e) 3’ overhang (multi-coloured cartoons), and negative controls (black cartoons) containing BITEG-modified 3’-ends. Results were normalised against their respective negative controls (absence of ExoIII) and converted to bp. The grey curve in (e) shows the equivalent reaction without an overhang, highlighting the loss of activity with a 4 bp 3’ overhang. **f**, Rate of resection by ExoIII on the blunt (red), 2-nt (orange) and 4-nt (green) 3’ overhangs and the negative control with BITEG-treated 3’-ends. Minimal loss of activity is observed with a 2-nt overhang, in contrast to almost complete loss of activity with a 4-nt overhang. **g**, Comparison of ExoIII and DNase I activity on their respective substrates. Error bars represent SEM; n=3 in all cases.

**Figure 3:**
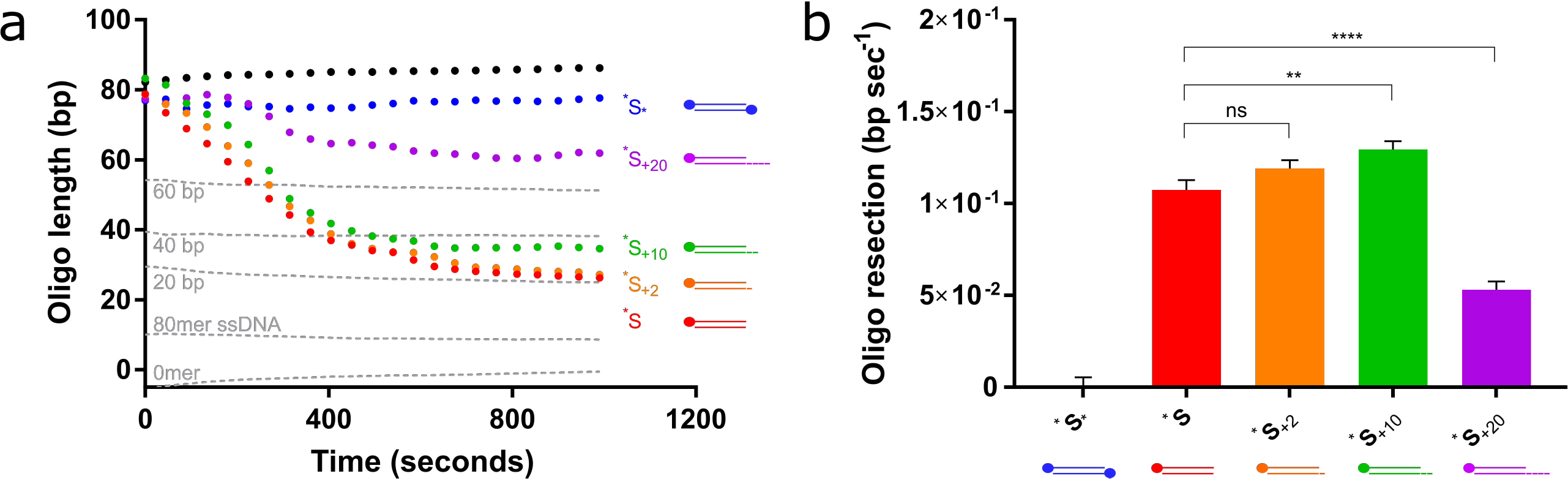
T7 Exo is inhibited by 20-nt 5’ overhangs. **a**, 12 nM T7 Exo-mediated resection on 50 nM substrates presenting a blunt (red), 2- (orange), 10- (green) and 20-nt (purple) 5’ overhang with. The negative control was synthesised with two 5’-BITEG modifications (blue). Results were normalised against their respective negative controls and converted to bp. **b,** Analysis of the rate of resection on the substrates in (a). Error bars represent SEM; n=3 in all cases; **p<0.01, ****p<0.0001.

The purpose of the standard curve is to enable the conversion of fluorescence units to DNA base pairs to calculate the length of substrate remaining following nucleolytic attack. To convert the data from fluorescence units, the data is first normalised to their relevant control (absence of enzyme). This has the additional benefit of accurately accounting for the inevitable photobleaching effect. As such, any decrease in fluorescence recorded is a result of nuclease activity and not photobleaching of the PG. We then convert the standard curve to base pairs by finding the gradient (m) and intercept (c) of the curve at each time point, and from this we can determine the length of dsDNA (x-axis) from the fluorescent signal (y-axis) at each time point and experimental condition. A worked example is available in Supplementary Materials.

We performed a DNase I titration and converted the y-axis from fluorescence units to length of dsDNA (Fig. 1d). From this we determined that the rate increases with the concentration of DNase I, therefore indicating that the reaction is first order with respect to the enzyme (Fig. 1e). We then calculated that the rate of digestion mediated by DNase I on a 2.5 ng/μL 80-bp dsDNA substrate at 37°C is approximately 0.001 bp nM^−1^ sec^−1^ (Fig. 1f).

### Optimisations for substrate-specific nucleases 3’-5’ nucleases: Exonuclease III

Oligonucleotides are far more versatile substrates than plasmids as they can be designed to have different terminal structures, such as 5’ or 3’ overhangs, Y-structures, Holliday junctions, strand invasion or even G-quadruplexes (44). Their much shorter length ensures that each enzyme is confronted with a higher proportion of these alternative structures. As such, only oligonucleotides were used for further development of the assay. Our oligonucleotides were designed and optimised so as to reduce the likelihood of secondary structure formation in the ssDNA (43).

To account for having a sufficient dsDNA region as well as structurally appropriate terminal ends to study nuclease structural preferences, we designed substrates that were blocked at one end in order to direct the enzyme to resect one single strand only, leaving a full-length, blocked, single-stranded 80-mer substrate as the end product. The substrates were blocked with biotin-TEG (BITEG) at the necessary terminal ends. BITEG alone is insufficient to inhibit resection on substrates blocked at both terminal ends. However, inclusion of streptavidin in the reaction mix successfully protected the modified ends from resection (Fig. S2a-b). We noticed that the presence of streptavidin, but not BITEG alone, is responsible for preventing total resection, prohibiting ExoIII from resecting the final 20 nucleotides (Fig. S2c). This combination of biotin and streptavidin has previously been shown to be effective (10). Substrates blocked at both ends while in the presence of streptavidin and treated with enzyme show 15-20% loss of fluorescence. The expected purity of the oligomers synthesised is approximately 85% and, as such, there is always a small proportion of substrate present that either lacks the BITEG modification, or is not synthesised to full-length, rendering them susceptible to resection (Fig. S2a-c). One additional benefit of using BITEG and streptavidin to block resection is that it does not require prior knowledge of inhibitory structures (e.g. designing 4-nt 3’-overhangs to inhibit ExoIII (45) or a 22-nt 5’-overhang specifically to inhibit T7 Exo (14)).

We ensured that the linear range is maintained with the addition of BITEG and streptavidin using a calibration curve (Fig. 2b). While the overall fluorescence is lower, the maximal usable concentration of substrate remains unchanged. We then tested whether the assay could be used to study the activity and structural preference of 3’-5’ exonucleases, such as ExoIII. ExoIII has been well-characterised, and so we attempted to replicate its known activities in our assay. A range of ExoIII titrations were performed on three different substrates: blunt ended, 2-nt 3’-overhang, and a 4-nt 3’-overhang. The substrates have been depicted as cartoons for simplicity, and have been named according to their modifications; for example, S^*^ indicates a 3’-terminal BITEG modification, and _+2_S^*^ indicates a 3’-BITEG modification, and a 2-nt 3’ overhang on the opposite strand. ^*^S would instead reference a 5’-BITEG modification. The preference of ExoIII for blunt and 2-nt overhangs is known, as well as the inhibitory effect of 4-nt 3’-extensions, as these activities are often exploited for various sequencing and DNA detection techniques (46–49) (Fig. 2c-e). After determining the rate of resection on all substrates (Fig. 2f-g), it is clear that this assay is readily able to detect the previously reported substrate specificity of ExoIII. ExoIII is completely inhibited by 4-nt 3’-overhangs, and resects with this with a similarly low rate as the negative control, which has both 3’-ends blocked with BITEG and streptavidin. There is a peak of activity at 10 nM ExoIII, in which the rate of resection appears to be twice as efficient compared to higher and lower concentrations of the enzyme.

Through analysis of the efficiency of each nuclease, we can also compare the activities of ExoIII and DNase I. These results suggest that the ExoIII resection activity is approximately five-fold faster than the DNase I nuclease activity (Fig. 2g) at 37°C in their respective buffers.

### 5’-3’ exonuclease: T7 exonuclease

Having demonstrated that this assay is suitable for 3’-5’ nucleases, we next explored whether the assay could be used to study 5’-3’ nucleases. This necessitated the synthesis of substrates presenting 5’-BITEG modifications. We tested the affinity of T7 Exo for a selection of substrates, incorporating blunt, 2-, 10- and 20-nt 5’-overhangs (Fig. 3a). T7 Exo shows no significant difference between the affinity for blunt and 2’-nt 5’-overhangs. As had previously been observed, the rate of T7 Exo-mediated resection with a 20-nt overhang was approximately 50% of a short overhang (14) (Fig. 3b). Unexpectedly, as the length of the overhang increased, resection was slightly delayed. However, after resection commenced, it appeared to take place at a faster rate on the intermediate overhangs, considering that the reaction culminates at approximately the same time point on all substrates (p<0.002 between ^*^S and ^*^S_+10_, and p<0.0001 between ^*^S and ^*^S_+20_) (Fig. 3b). Regardless, difference in rate is rather minimal. In contrast to ExoIII, T7 Exo is limited, but not inhibited, by the 20-nt overhang.

### Studying enzyme resection through single-nucleotide mismatches

During mismatch repair, DNA nucleases resect through the mismatch to generate a single-stranded tract of DNA along which DNA polymerases can replicate (50–54). It has been suggested that these resection events may even remove epigenetic signatures on the DNA, such as 5-methylcytosine (55), which may then be permanently lost when repaired with an unmodified cytosine, and not targeted by *de novo* methyltransferases, Dnmt3a or Dnmt3b (56, 57). We therefore tested if our assay can detect differences in resection rates through mismatched and methylated substrates.

We investigated ExoIII activity on mismatched substrates compared to matching substrates and identified some unexpected activities. First, we observed that ExoIII resects the two perfectly matching substrates at substantially different rates, resecting _*_S at a 1.7 - 2.0-fold faster rate than S^*^ (Fig. 4a-d) _*_S^C>T^ contains a selection of C>T substitutions, whilst _G>A_S^*^ contains G>A substitutions, generating T:G and A:C mismatches, respectively. The presence of the T:G mismatches slows the rate of resection 0.09 − 0.16X compared to its perfectly matching counterpart (p<0.001 for 5 nM ExoII; p<0.05 for 10 nM ExoIII; Fig 4d). Conversely, incorporating A:C mismatches increases the rate of resection by 1.5 – 1.8X (p<0.0001 with both 5 and 10 nM ExoIII). There is some evidence for a nucleotide preference for ExoIII, although this has not been repeated to the best of our knowledge (58). The rate of resection on both the perfectly matching substrate appears to have a delayed start, particularly for S^*^. Interestingly, ^*^S is then resected at an accelerated rate compared to both mismatched substrates. The mismatches resect at a slower rate but, as their resection begins at the offset, the reaction on the mismatched substrates and *S finish at the same time point for both 5 and 10 nM ExoIII.

**Figure 4:**
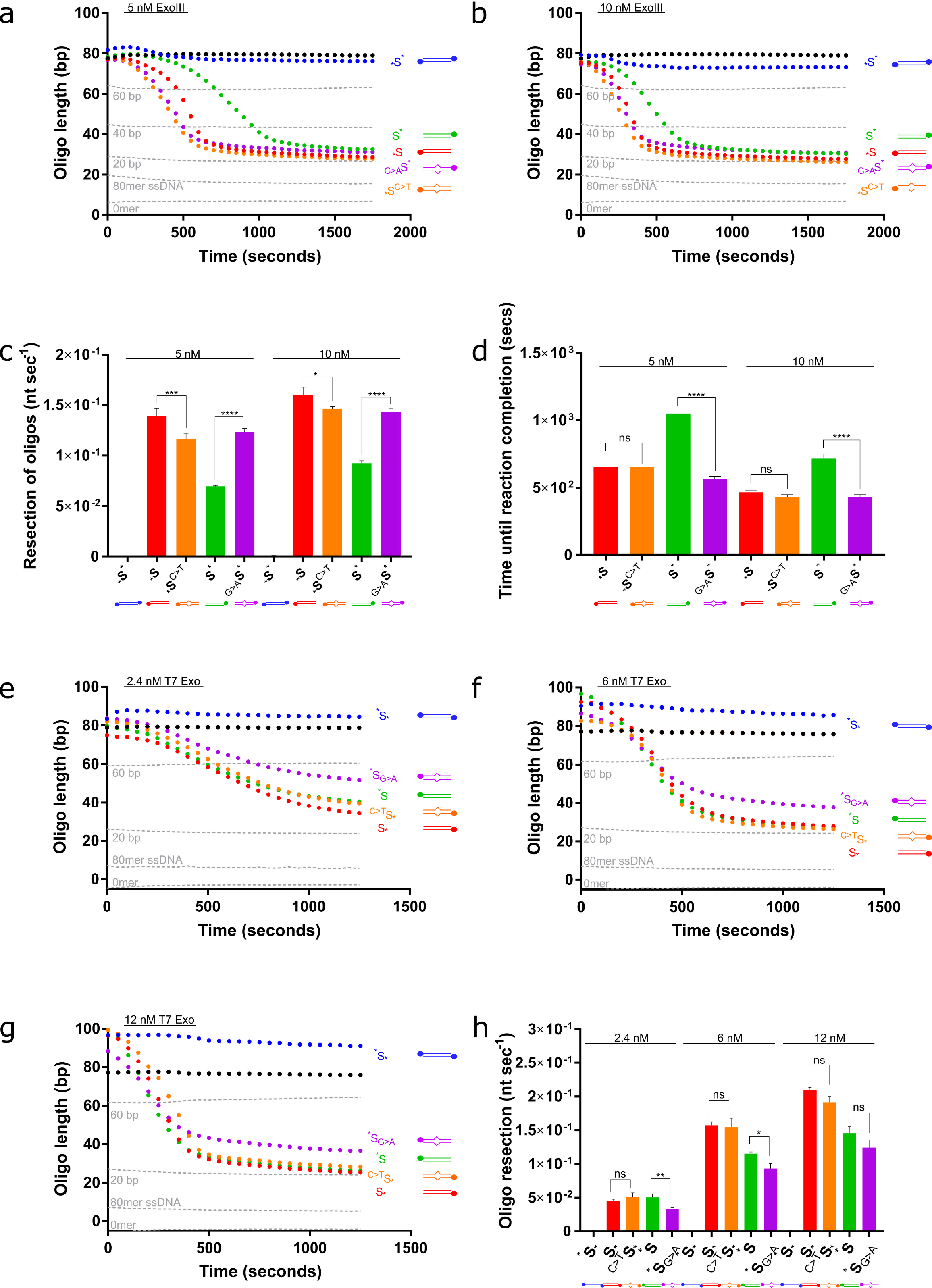
ExoIII and T7 Exo resect through single-nucleotide mismatches. **a,** 5 nM and **b,** 10 nM ExoIII was added to perfectly matched substrates (_*_S and S^*^) and substrates containing four T:G or A:C mismatches, (_*_S^C>T^ and _G>A_S^*^, respectively), and the resection reaction followed. Standard curve is represented by the grey dotted lines. **c,** Calculated resection rate of ExoIII on the complementary and mismatched substrates based on maximum gradients in (a) and (b). **d,** Time (seconds) until the ExoIII reaction reaches completion based on the point at which the reaction plateaus. **e,** 2 nM, **f,** 7.5 nM and **g,** 10 nM T7 Exo was added to complementary substrates (S_*_ and ^*^S) and substrates containing four T:G or A:C mismatches (^C>T^S_*_ and ^*^S_G>A_, respectively). Standard curve is represented by the grey dotted lines. **h,** Calculated resection rate of T7 Exo on the complementary and mismatched substrates. Error bars represent SEM; n=3 in all cases; *p<0.05, **p<0.01, ***p<0.001, ****p<0.0001.

T7 exonuclease is considered to be sequence-independent as it lacks a defined DNA binding-motif, similar to other FEN family nucleases that bind DNA in a nonspecific manner (14, 59). Nevertheless, it has previously been suggested that it might resect different nucleotides with variable efficiency (60). Indeed, we observed that T7 Exo resects the perfectly complementary ^*^S with more difficulty than S_*_, which was noticeable at higher concentrations of T7 Exo (Fig. 4e-g). Similarly with T7 Exo, our assay detected a substrate strand preference (Fig. 4h).

The addition of four T:G mismatches did not slow or accelerate T7 Exo activity. However, incorporating A:C mismatches into the more resistant substrate did appear to slow resection, and this general trend was observed across all concentrations of T7 Exo. The effect of the mismatches was more evident at lower T7 Exo concentrations (p<0.05 for 7.5 nM and p<0.01 for 2 nM T7 Exo). These data suggest that mismatches may only be inhibitory to T7 Exo in certain sequence contexts. Indeed, T7 Exo has been shown to recognise single-nucleotide mismatches in an SNP-detection system (61).

### Enzyme resection through methylated cytosines

Neither ExoIII nor T7 Exo is substantially inhibited by methylated DNA in this experimental system. At the lower concentration of ExoIII, we see a delay in resection initiation (Fig. 5a), resulting in staggered end points (Fig. 5c). However, once ExoIII starts to resect, the rates are not significantly different between the methylated and non-methylated substrates (Fig. 5d). This delay in resection is not observed at higher concentrations of ExoIII (Fig. 5b). T7 Exo showed no significant effect on either maximum rate or time taken to resect in the presence of methylated cytosines (Fig. 5e-h). To our knowledge, there is no data on the effect of methylcytosine on either ExoIII or T7 Exo resection with which to compare.

**Figure 5:**
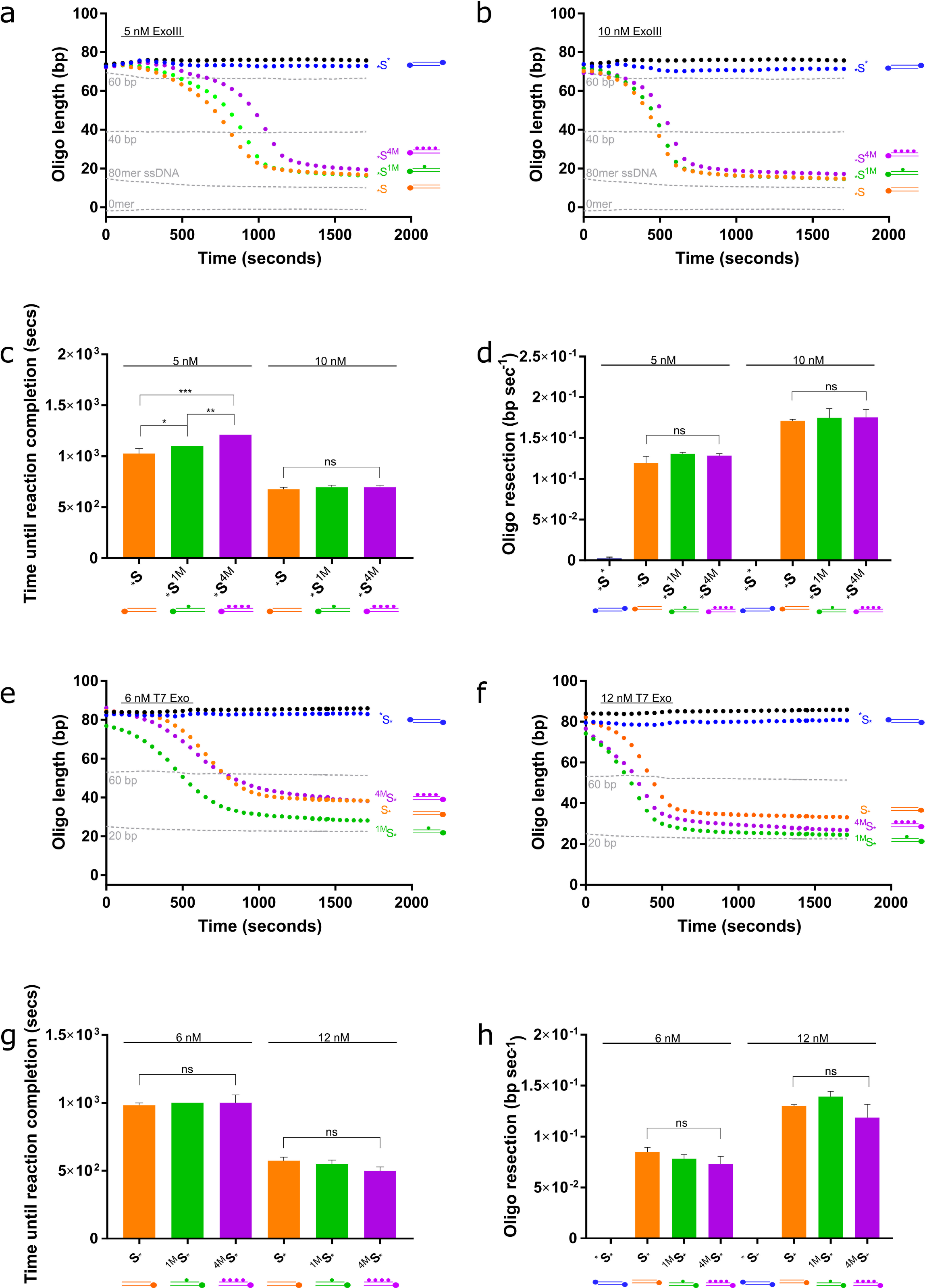
Increased methylcytosine content delays ExoIII-mediated resection; but does not affect the rate of resection of either ExoIII or T7 Exonuclease. **a,** 5 nM and **b,** 10 nM ExoIII was added to a non-methylated substrate, a substrate containing one methylated cytosine, and a substrate containing four methylated cytosines (*S, _*_S^1M^ and _*_S^4M^, respectively). Standard curve is represented by the grey dotted lines. **c,**Time (seconds) until the ExoIII reaction reaches completion on the methylated and unmethylated substrates based on the point at which the graphs plateau in (a) and (b). **d,** Calculated resection rate of ExoIII based on maximum gradients in (a) and (b). **e,** 6 nM and **f,** 12 nM T7 Exo on non-methylated and differentially methylated substrates. Standard curve is represented by the grey dotted lines. **g,** Time (seconds) until the T7 Exo reaction reaches completion on the methylated and unmethylated substrates based on point at which the graphs plateau in (e) and (f). **h,** Calculated resection rate of ExoIII. Error bars represent SEM; n=3 in all cases; *p<0.05, **p<0.01, ***p<0.001.

### Validation of nuclease activity assay on nicks and gaps, and its use in studying nickases in combination with processive nucleases

Nicks and gaps are introduced as intermediates in DNA repair mechanisms, yet they need to be rapidly processed to prevent the accretion of double-strand breaks at replication forks, possibly leading to cell death (10, 62).

Due to the physiological relevance of nicks and gaps, we explored the nucleolytic activity of ExoIII and T7 Exo on these structures. In order to test this, we used a combination of substrates that had been designed to contain a nick or gap, in conjunction with blocking at both ends (both 5’ or 3’, respectively) to prevent resection from the terminal ends. We also included a recently-purified nickase, Nt.CviPII (41) which is known to possess inherent exonuclease activity. Nickases cleave just one strand of duplex DNA, breaking the phosphodiester backbone. Nt.CviPII preferentially cuts CCA and CCG, but cuts less efficiently at CCT (41). One strand of the dsDNA substrate contains five evenly distributed CCA motifs, while the opposite strand only contains two CCT motifs, and therefore most of ExoIII’s activity should be directed on the first strand (Fig. 6a). To minimise the exonuclease activity of Nt.CviPII, we used a low dilution of the nickase.

**Figure 6:**
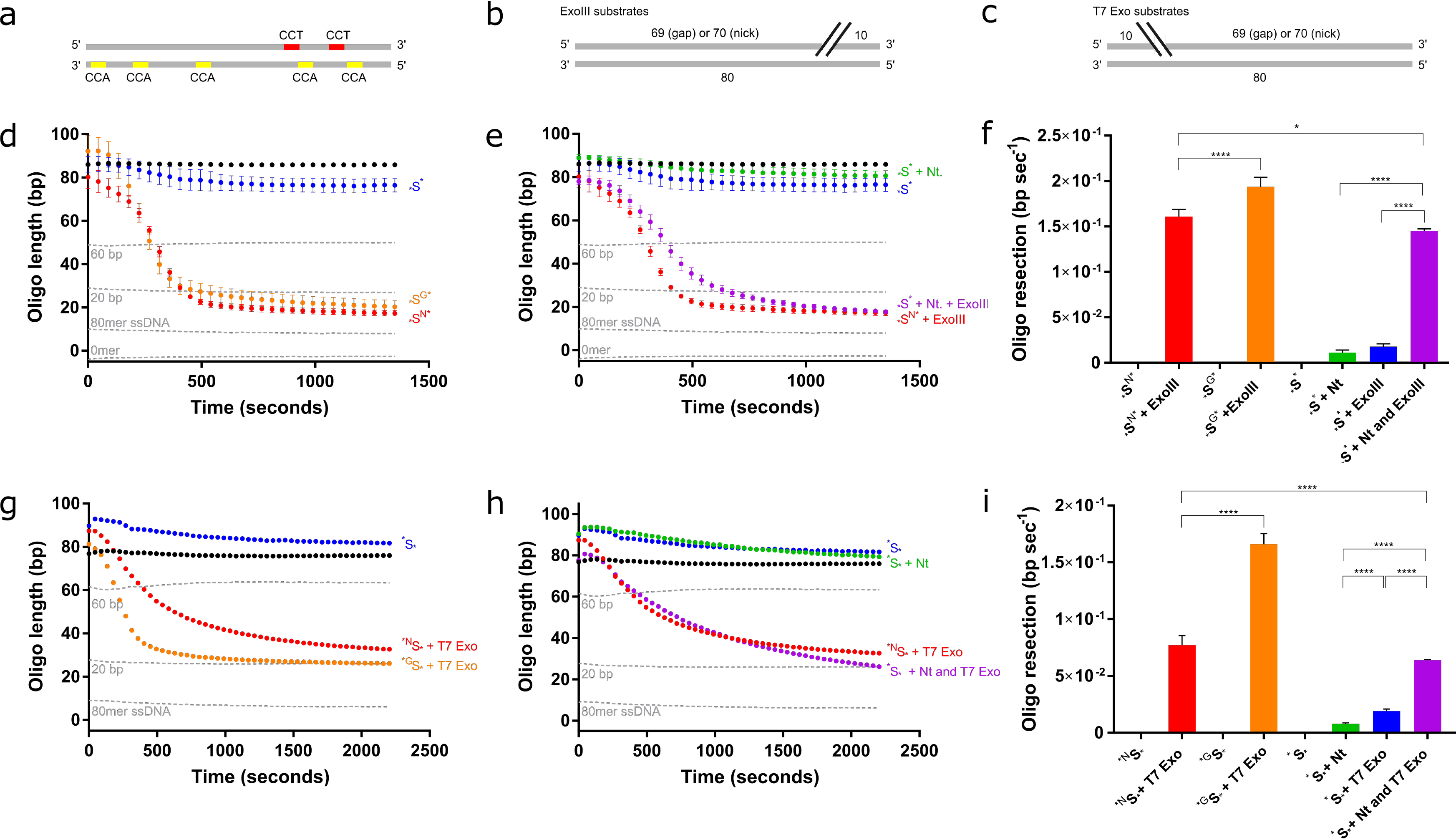
ExoIII and T7 Exo preferentially resect from a nick than a gap. **a**, Schematic of the dsDNA substrate indicating the preferred nicking sites (yellow) and the least favourable nicking sites (red) of Nt.CviPII. **b-c,** Schematic of the dsDNA substrates presenting a nick or gap at either the 3’-end for ExoIII, or the 5’-end for T7 Exo. **d,** ExoIII is active against simulated nicked substrates. 40 nM ExoIII was added to a blocked substrate (blue) and related substrates designed with either a nick (red) or a gap (orange) towards the 3’ end of one strand, showing similar activity against both modified substrates. Standard curve is represented by the grey dotted lines. **e,** Quantification of the Nt.CviPII nickase enzyme activity coupled to ExoIII. Nt.CviPII was added to the blocked substrate, either with (purple) or without (green) 40 nM ExoIII. Controls from b) are shown for comparison. **f,** Calculated resection rate of nicked and gapped substrates by ExoIII, extracted from the maximum gradient. Addition of the nickase significantly increases the resection rate, highlighting that the nickase activity is detected. **g,** T7 Exo is active against simulated nicked substrates. 12 nM T7 Exo was added to substrates with a nick (red) or a gap (orange) towards the 5’ end of one strand. Negative control is in blue. Standard curve is represented by the grey dotted lines. **h,** Quantification of the Nt.CviPII enzyme activity coupled to T7 Exo. Nt.CviPII was added to the blocked substrate, either with (purple) or without (green) 12 nM T7. Negative control is in blue. **i,** Calculated resection rate of nicked and gapped substrates by T7 Exo, extracted from the maximum gradients in (g) and (h). Addition of the nickase significantly increases the resection rate, highlighting that the nickase activity is detected. Error bars represent SEM; n=3 in all cases; *p<0.05, ****p<0.0001.

Our results show that ExoIII functions marginally better on gaps rather than nicks (p<0.0001) (Fig. 6d and 6f), and this may reflect its role in base excision repair where it resects from an abasic site (39). ExoIII is nevertheless able to resect from a nick, and the rate is comparable to its activity on blunt ends, as previously observed (63). ExoIII also resected from the nickase-induced nicks at an almost equal rate to the substrates designed to contain a single nick (Fig. 6e and 6f). These data demonstrate that Nt.CviPII is a very fast acting nickase against its preferred substrate sequences as there is no delay in the start of ExoIII-mediated resection.

As with ExoIII, T7 Exo also resects from both nicks and gaps. It has a more defined preference for gaps, showing a rate approximately 2.5-fold greater than for nicks (p<0.0001; Fig. 6g and 6i). T7 Exo also resects from nicks generated by a nickase almost as efficiently as from a substrate already presenting a single nick (Fig. 6h and 6i). Typically, T7 endonuclease cleaves at nicked sites during infection to generate DNA double-stranded breaks that are susceptible to T7 Exo. As such, it is perhaps a redundant property of T7 Exo to resect from a nick (64).

### Alternative enzymes that degrade or synthesise dsDNA can be studied

We have demonstrated that PG is an effective dye that can be used to study dsDNA nucleases from viruses and bacteria. Given the low, but not insignificant, fluorescent signal observed for the single-stranded 80-mer control in the standard curves, we reasoned that this assay may have the sensitivity to address ssDNA nucleases, such as human Trex2 (65). We confirmed this by showing that the assay is sensitive enough to detect a Trex2 concentration-dependent decrease in fluorescence from ssDNA substrates ranging in size from 60 to 20 nts (Fig.7a). This indicates that this assay has considerable potential for an even wider range of DNA nucleases, and very high levels of sensitivity in order to capture such activity in detail. As expected, it takes longer for Trex2 to digest longer substrates (Fig. 7b). In terms of reaction kinetics, rate of resection is generally consistent, ranging between 1.7 x 10^−4^ − 3.4 x 10^−4^ nt nM^−1^ sec^−1^ (Fig. 7c), which is comparable to previously published rates (66).

**Figure 7:**
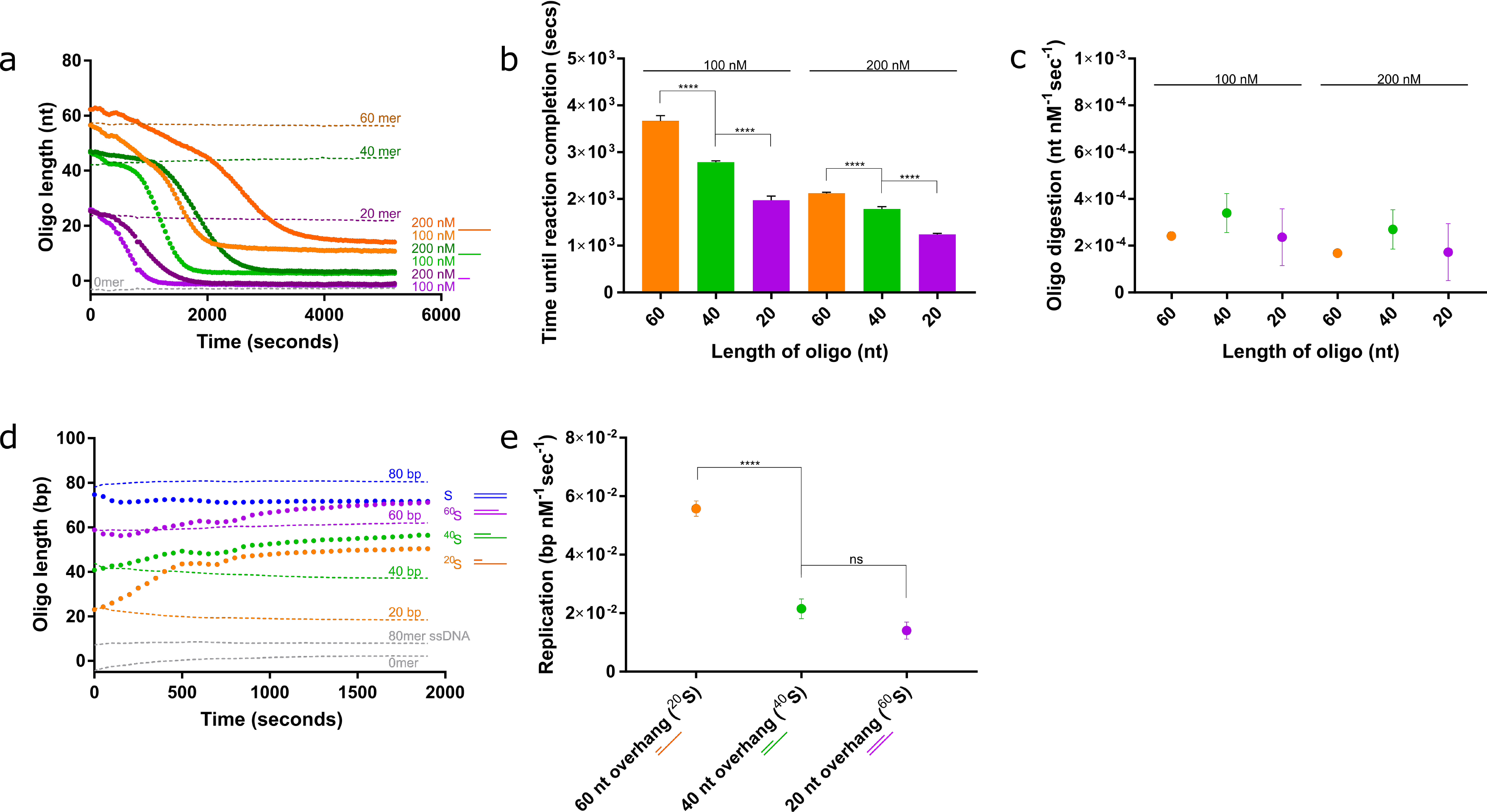
Validation of the assay for alternative enzymes that digest ssDNA and generate dsDNA. **a,** Mammalian Trex2 digestion of single stranded DNA quantified using the PG assay. Single-stranded 60-mer (orange), 40-mer (green) and 20-mer (purple) DNA substrates were treated with 100 and 200 nM Trex2. Robust digestion was observed in each case. **b,** Time (seconds) until reaction completion based on the point at which the graphs plateau in (a). **c,** Rate of digestion per nM Trex2. Trex2 degrades DNA at a similar rate irrespective of the length of the DNA substrate. **d,** The polymerase activity of the Klenow fragment polymerase determined using the PG assay. Three substrates with a 20-, 40- and 60- and overhang with a total 80 bases were incubated with 1 nM Klenow fragment. **e,** Calculated polymerisation rate of the Klenow fragment. A clear preference is shown for longer overhangs. Error bars represent SEM; n=3 in all cases; ****p<0.0001.

Since we have shown that this assay is a powerful tool for observing nuclease activity, we attempted to study whether we could capture the reverse activity and visualise an increase in fluorescence upon addition of polymerase. To investigate this, we employed the use of the well-characterised Klenow fragment polymerase (KF). KF requires a short DNA primer fragment hybridised to a ssDNA fragment along which it can replicate. We used a selection of 80-mer oligomers hybridised to a 20-, 40-, 60- and 80-mer oligomer (Fig. 7d). The data clearly indicate that KF-dependent blunting of the 20mer 5’-overhang is possible to visualise in the context of this real-time assay. KF appears to be unable to extend the 20- and 40-mer primers to produce the full-length 80 bp dsDNA product. KF-elongation of both are inhibited once the primers have been extended to 50- or 55-mer lengths (Fig. 7d). It is understood that KF is sensitive to secondary structures in the ssDNA template, and the presence of a small hairpin adjacent to this region may be responsible for inhibiting polymerisation past this point. Previously, a terminal hairpin has not inhibited KF processivity, although an internal hairpin, such as in this case, may have a different impact on KF (67). Analysis of the rate of replication suggests a preference for a longer ssDNA template, as KF polymerises at a much faster rate on the 60mer 5’-overhang, while the 40- and 20-mer 5’-overhangs are processed at a three-fold lower rate (Fig. 7e).

### RNA and RNA:DNA nucleases can be also be analysed in real-time

We have established a straightforward and robust assay that can measure DNA nuclease activity. To expand the potential of this assay to include RNA nucleases, we employed the use of an alternative RNA-sensitive fluorescent dye available in the Quanit-iT microRNA assay kit, as PG emits very low fluorescent signal in the presence of RNA.

With the use of RNA nucleases, RNase A and RNase H, we ascertained that we could reproduce their known activities. Indeed, we observed that RNase A efficiently resected ssRNA (Fig. 8a), at a rate of approximately 180 nt nM^−1^ sec^−1^ on a 20-nt substrate, and 90 nt nM sec^−1^ on a 40-nt substrate (Fig. 8b). As expected, RNase A failed to show any nucleolytic activity on dsRNA, RNA:DNA hybrids, or dsDNA (Fig. S3a-c). Interestingly, in this buffer the 20- and 40-mer RNA substrates emitted very similar fluorescent signals and could not be teased apart, yet the fluorescent signals were easily distinguishable on dsRNA and RNA:DNA hybrid substrates (Fig. S3a-b).

**Figure 8:**
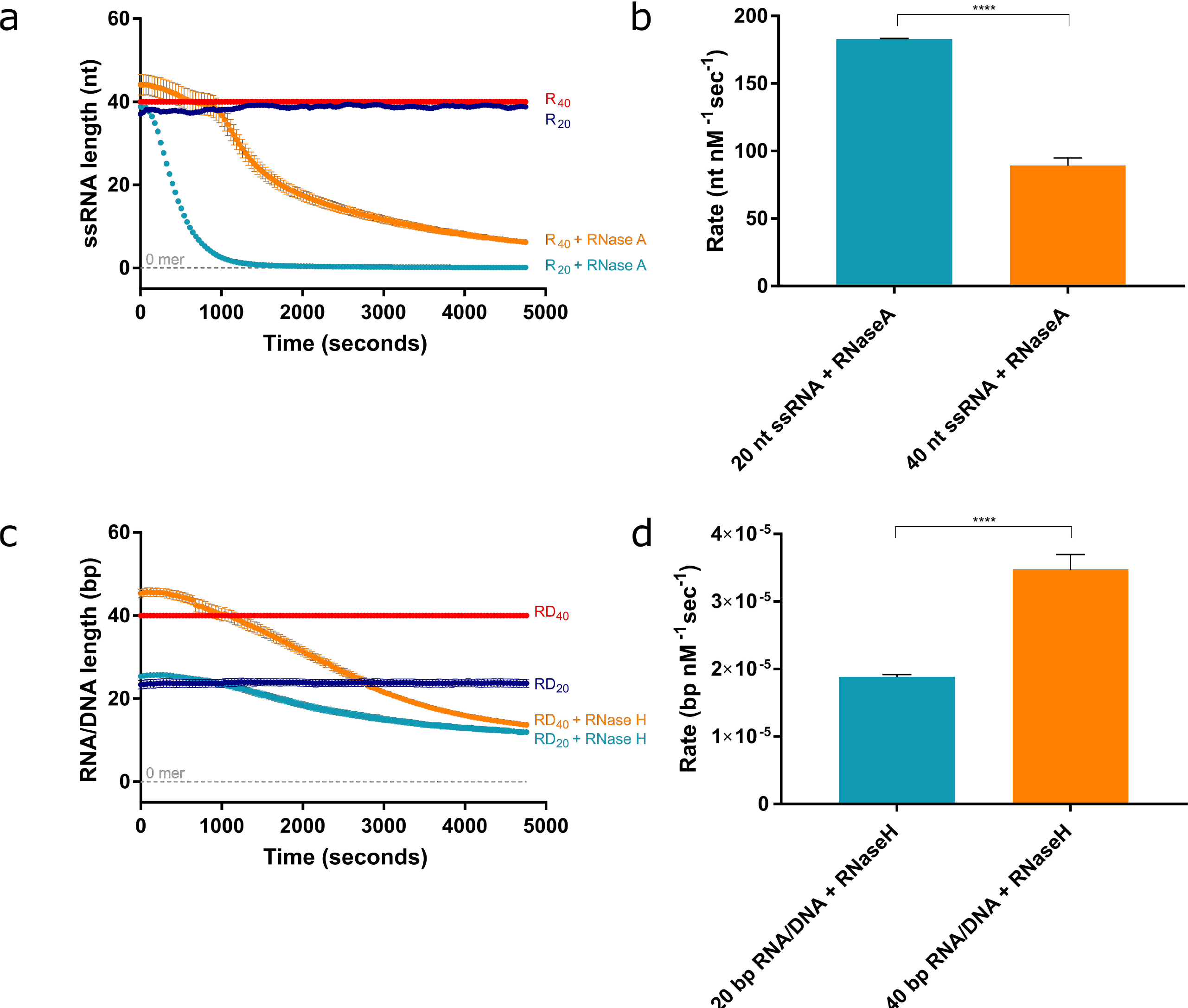
QI fluorescent-based toolkit reiterates RNase A and RNase H activities ssRNA and DNA/RNA hybrids, respectively. **a,** Robust digestion of 20- and 40-mer ssRNA by 3.64 x 10^−4^ nM RNase A. **b,** Quantification of the rate of digestion according to the maximum gradient in (a). **c,** Robust resection of the RNA strand of the 20- and 40-mer RNA:DNA hybrids by 280 nM RNase H. **d,** Quantification of the rate of resection based on the maximum gradient in (c). Error bars represent SEM; n=3 in all cases; ***p<0.001.

On the other hand, RNase H exerts a distributive cleavage pattern on the RNA substrate of RNA:DNA hybrids, periodically cleaving the RNA to produce fragments (Fig. 8c). RNase H did not show any affinity for ssRNA, dsRNA, or dsDNA substrates (Fig. S4a-c). This may account for the much slower rates exhibited by RNase H, as it digested the RNA component of RNA:DNA hybrid substrates at 2.0 x 10^−5^ nt nM^−1^ sec^−1^ on the 20-bp substrate, and 3.5 x 10^−5^ nt nM^−1^ sec^−1^ on the 40-bp substrate (Fig. 8d).

## Discussion

We have further developed a highly-sensitive, fluorescence-based universal nuclease assay that works in real-time and designed a library of substrates that are appropriate for investigating a wide range number of nucleases, polymerases, and helicases. This assay can be used to calculate reaction kinetics and reaction completion times, providing a powerful quantitative tool for characterising enzymes active on nucleic acids. We have successfully validated the technique using well-studied enzymes and, in the process, identified previously unreported details of their mechanisms.

Our assay has important applications in comparing the relative activities of enzymes. Here, we identified that ExoIII hydrolyses the DNA substrate at a faster rate than DNase I in their respective buffers. DNase I exhibits a relatively low affinity for DNA, and higher affinity strains have been engineered for treatment of cystic fibrosis (68, 69). DNase I may have evolved to be less efficient due to the fact that it can digest all structures of DNA, irrespective of whether it is single- or double-stranded, and deregulation of this activity could be disastrous for the cell. This limitation in its binding may exert some control to prevent inappropriate activity from causing cellular damage. ExoIII shows a greater rate of resection per enzyme protomer, yet this is counteracted by its very limited structural specificity. The product of ExoIII resection during *E. coli* base excision repair is a single-stranded tract of DNA that can be much more easily repaired than the damage caused by DNase I (70, 71).

We were also able to compare substrate specificities, as indicated by the preference of ExoIII for 3’-overhangs shorter than 4-nt and T7 Exo’s predilection for 5’-overhangs of 10-nt or fewer. As T7 Exo typically resects DNA from short 5’-overhangs introduced by T7 endonuclease during the infection process in *E. coli*, our results are consistent with its behaviour *in vitro* and *in vivo* (14, 64). In addition to substrate structures, we have considered other physiologically relevant substrates containing single-nucleotide mismatches and methylcytosines. Unexpectedly, we found that ExoIII and T7 Exo exhibited a preference for one strand of the DNA substrate. There is evidence of a nucleotide preference for ExoIII; C>A~T>G, although it seems that it depends on the sequence context (58). Nevertheless, the results of their mismatched counterparts do coincide with published ExoIII preferences (58). As such, this confirms previously reported preferences in both a qualitative and quantitative way. Work is continuing in our lab to understand the mechanisms of ExoIII, and so we hope to elucidate its nucleotide preferences thoroughly.

For methylated substrates, we observed that resection is delayed for lower concentrations of ExoIII and are inhibited slightly by the presence of multiple methylcytosines. The rate of resection, however, remains consistent on all substrates. At higher concentrations, ExoIII does not appear to distinguish the methylcytosines. T7 Exo does not show any alteration in activity from the methyl groups. Its host, *E. coli*, contains a small percentage of methylated adenines and cytosines, and it appears that T7 Exo does not discriminate against them. This is in spite of a smaller percentage of methylated DNA ending up in the T7 bacteriophage’s progeny DNA than is present in the *E. coli* genome (72).

While this assay is predominantly useful for studying processive enzymes, it is possible to combine a nickase with an enzyme that resects from nicks, as indicated with nickase Nt.CviPII in conjunction with either ExoIII or T7 Exo. This is a powerful method to identify nicking or endonuclease activities, and study their efficiency based on when resection commences.

The superior sensitivity of PG in this assay also expanded our repertoire of enzymes to include ssDNA nucleases, such as Trex2. Trex2 is one of at least eight autonomous exonucleases in human cells, and is likely recruited to 3’-termini to alleviate blocks during replication arrest (73). It binds DNA very tightly, and this affinity for its substrate has been captured here due to its very rapid processivity. In addition to the study of enzymes that digest DNA, we extended our study to enzymes that polymerise DNA, such as the nuclease-deficient Klenow fragment polymerase. Its polymerase activity is ten-fold more rapid, for example, than ExoIII at resection in their respective buffers

We further explored the fluorescent dye QI which, unlike PG, recognises RNA polynucleotides. The activities of RNase A and H were captured in real-time, where we recapitulated their known activities on ssRNA and the RNA strand of RNA:DNA heteroduplexes, respectively.

Characterisation of DNA nucleases is integral for demystifying their roles in maintenance of genomic integrity. Loss of nucleases, or mutations in their structural or functional domains, can have disastrous effects on the health of an organism. The impact of these mutations could affect resection rates or binding affinity to the DNA or interacting partners, and we posit that this assay may be sensitive enough to capture and compare these characteristics (69, 74, 75). Many nucleases have been well-characterised, including those used in the optimisation of this assay, while many more have yet to be characterised at all.

Nucleases may also represent anticancer targets, and this assay could offer an alternative method for studying the effect of future anticancer drugs on the activity of their target nuclease (76). In a similar instance, a discontinuous assay using PG was successfully performed to study the inhibitory role of actin on DNase I (77). As such, this assay represents a safe, easy, rapid, robust, real-time study of dsDNA and ssDNA nucleases and polymerases. We believe that it has the potential to revolutionise quantitative assessment of DNA and/or RNA cleaving enzymes in a vast range of applications.

## Additional information

See attached supplementary information

## Acknowledgements and Funding

We are grateful to Dr Mirella Vivoli (University of Florence) for her insight in the early development of this assay; and to Fulvia Bono (University of Exeter) for her insight on RNA nucleases and her support of this project. We are also very thankful to Adam Thomas (University of Exeter) for his knowledge and guidance on the data analysis. We also thank Matthew Scharff and Shanzhi Wang (Albert Einstein College of Medicine), and Michael Dillon (University of Exeter) for helpful suggestions. Funding for RC was provided by Biotechnology and Biological Research Council [BB/N017773/1], Royal Society [IE150290], and Academy of Medical Sciences [SBF001\1005]. ECS is funded by a PhD studentship from the Biotechnology and Biological Research Council-funded South West Doctoral Training Partnership [BB/J014400/1].

## Competing Interests

The authors declare no competing interests.

